# Dysregulation of the Tweak/Fn14 pathway in skeletal muscle of spinal muscular atrophy mice

**DOI:** 10.1101/2021.09.13.460053

**Authors:** Katharina E. Meijboom, Emily McFall, Daniel Anthony, Benjamin Edwards, Sabrina Kubinski, Gareth Hazell, Nina Ahlskog, Peter Claus, Kay E. Davies, Rashmi Kothary, Matthew J.A. Wood, Melissa Bowerman

## Abstract

Spinal muscular atrophy (SMA) is a childhood neuromuscular disorder caused by depletion of the survival motor neuron (SMN) protein. SMA is characterized by the selective death of spinal cord motor neurons, leading to progressive muscle wasting. Loss of skeletal muscle in SMA is a combination of denervation-induced muscle atrophy and intrinsic muscle pathologies. Elucidation of the pathways involved is essential to identify the key molecules that contribute to and sustain muscle pathology. The tumor necrosis factor-like weak inducer of apoptosis (TWEAK)/TNF receptor superfamily member fibroblast growth factor inducible 14 (Fn14) pathway has been shown to play a critical role in the regulation of denervation-induced muscle atrophy as well as muscle proliferation, differentiation and metabolism in adults. However, it is not clear whether this pathway would be important in highly dynamic and developing muscle. We thus investigated the potential role of the TWEAK/Fn14 pathway in SMA muscle pathology, using the severe Taiwanese *Smn^-/-^;SMN2* and the less severe *Smn^2B/-^* SMA mice, which undergo a progressive neuromuscular decline in the first three post-natal weeks. Here, we report significantly dysregulated expression of the TWEAK/Fn14 pathway during disease progression in skeletal muscle of the two SMA mouse models. In addition, siRNA-mediated *Smn* knockdown in C2C12 myoblasts suggests a genetic interaction between Smn and the TWEAK/Fn14 pathway. Further analyses of SMA, *Tweak^-/-^* and *Fn14^-/-^* mice revealed dysregulated myopathy, myogenesis and glucose metabolism pathways as a common skeletal muscle feature, and providing further evidence in support of a relationship between the TWEAK/Fn14 pathway and Smn. Finally, a pharmacological intervention (Fc-TWEAK) to upregulate the activity of the TWEAK/Fn14 pathway improved disease phenotypes in the two SMA mouse models. Our study provides novel mechanistic insights into the molecular players that contribute to muscle pathology in SMA and into the role of the TWEAK/Fn14 pathway in developing muscle.

## BACKGROUND

The neuromuscular disease spinal muscular atrophy (SMA) is the leading genetic cause of infant mortality [1]. SMA is caused by mutations in the *survival motor neuron 1* (*SMN1*) gene [2]. The major pathological components of SMA pathogenesis are the selective loss of spinal cord alpha motor neurons and muscle wasting [3]. Skeletal muscle pathology is a clear contributor to SMA disease manifestation and progression and is caused by both denervation-induced muscle atrophy [4, 5] and intrinsic defects [6–8]. As skeletal muscle is the largest insulin-sensitive tissue in the body and is involved in glucose utilization [9], it is not surprising that muscle metabolism is also affected in SMA. Impaired metabolism has indeed been reported in SMA Type 1, 2 and 3 patients [10–14]. A better understanding of the specific molecular effectors that contribute to SMA muscle physiopathology could provide mechanistic insights in SMA muscle pathology and help therapeutic endeavors aimed at improving muscle health in patients [15].

One pathway that plays a crucial role in chronic injury and muscle diseases is the tumor necrosis factor-like weak inducer of apoptosis (TWEAK) and its main signaling receptor, the TNF receptor superfamily member fibroblast growth factor inducible 14 (Fn14) [16–18]. TWEAK is ubiquitously expressed and synthesized as a Type II transmembrane protein but can also be cleaved by proteolytic processing and secreted as a soluble cytokine [19]. The role of the TWEAK/Fn14 pathway in skeletal muscle is conflicting as it has been demonstrated to have both beneficial and detrimental effects on muscle health and function [20, 21]. Indeed, pathologically high levels of TWEAK activate the canonical nuclear factor kappa-light-chain-enhancer of activated B cells (NF-κB) pathway, which promotes myoblast proliferation and thus inhibits myogenesis and the early phases of muscle repair and regeneration [22, 23]. Conversely, lower physiological concentrations of TWEAK activate the non-canonical NF-κB pathway that promotes myoblast fusion and myogenesis [24]. The transmembrane protein Fn14 is typically dormant or present in low levels in normal healthy muscle [25]. Atrophic inducing conditions (e.g. casting and surgical denervation) stimulate the expression of Fn14, leading to the chronic activation of the TWEAK/Fn14 pathway and sustained skeletal muscle atrophy [26].

We have also demonstrated an increased activity of the Tweak/Fn14 pathway in skeletal muscle of a mouse model of the neurodegenerative adult disorder amyotrophic lateral sclerosis (ALS), which is characterized by a progressive and chronic denervation-induced muscle atrophy [27]. In addition, various downstream effectors of the TWEAK/Fn14 pathway play critical roles in the regulation of muscle metabolism such as peroxisome proliferator-activated receptor-gamma coactivator 1α (PGC-1α), glucose transporter 4 (Glut-4), myogenic transcription factor 2d (Mef2d), hexokinase II (HKII) and Krüppel-like factor 15 (Klf15) [28–34].

Although the TWEAK/Fn14 pathway has been ascribed roles in both skeletal muscle health regulation and metabolism, both of which are impacted in SMA [35, 36], this pathway has yet to be investigated in the context of SMA. Furthermore, all research on this pathway has been performed in adult mice and therefore has never been explored in early phases of muscle development. We thus investigated the potential role of TWEAK/Fn14 signaling in SMA and in early phases of post-natal skeletal muscle development. We report significantly decreased levels of both *Tweak* and *Fn14* during disease progression in two distinct SMA mouse models (*Smn^-/-^;SMN2* and *Smn^2B/-^*) [37, 38]. We also observed dysregulated expression of *PGC-1α*, *Glut-4*, *Mef2d* and *HKII*, the metabolic downstream effectors of TWEAK/Fn14 signaling [29, 30], in skeletal muscle of these SMA mice. In addition, more in-depth analyses revealed an overlap of aberrantly expressed genes that regulate myopathy, myogenesis and glucose metabolism pathways in skeletal muscle of SMA, *Tweak^-/-^* and *Fn14^-/-^* mice, further supporting shared functions between the TWEAK/Fn14 pathway and SMN in developing muscle. Finally, upregulation of the activity of the TWEAK/Fn14 pathway, through a pharmacological intervention (Fc-TWEAK administration), improved disease phenotypes in the two SMA mouse models. Our study uncovers novel mechanistic insights into the molecular effectors that contribute to skeletal muscle pathology in SMA and demonstrates a role for the TWEAK/Fn14 pathway in the early stages of post-natal muscle development.

## METHODS

### Animals and animal procedures

Wild-type mice FVB/N [39] and C57BL/6J [40] and the severe *Smn^-/-^*;*SMN2* mouse model (FVB.Cg-Smn1tm1Hung Tg(SMN2)2Hung/J) [41] were obtained from Jackson Laboratories. The *Smn^2B/-^* mouse model [38, 42] was kindly provided by Dr. Lyndsay M Murray (University of Edinburgh). *Tweak^-/-^* [43] and *Fn14^-/-^* mouse models [44] were generously obtained from Linda C. Burkly (Biogen).

Most experiments with live animals were performed at the Biomedical Services Building, University of Oxford. Experimental procedures were authorized and approved by the University of Oxford ethics committee and UK Home Office (current project license PDFEDC6F0, previous project license 30/2907) in accordance with the Animals (Scientific Procedures) Act 1986. Experiments with the *Smn^2B/-^* mice in Figure 1 were performed at the University of Ottawa Animal Facility according to procedures authorized by the Canadian Council on Animal Care.

**Figure 1.**
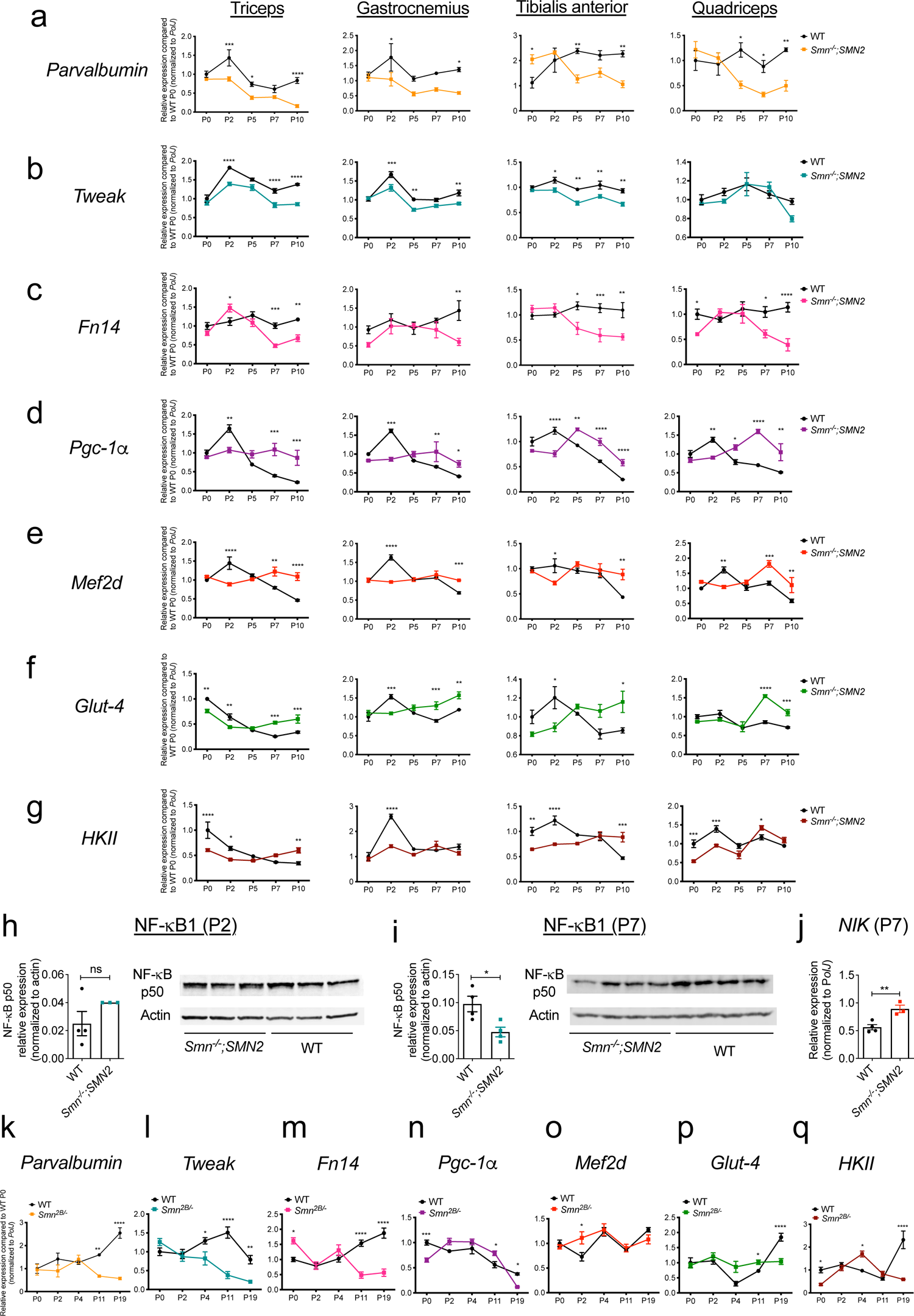
Aberrant expression of the TWEAK/Fn14 signaling pathway in skeletal muscle of SMA mice. **a-g**. qPCR analysis of *parvalbumin* (**a**), *Tweak* (**b**), *Fn14* (**c**), *Pgc-1α* (**d**), *Mef2d* (**e**), *Glut-4* (**f**) and *HKII* (**g**) in triceps, gastrocnemiu*s*, TA and quadriceps muscles from post-natal day (P) 0 (birth), P2 (pre-symptomatic), P5 (early-symptomatic), P7 (late symptomatic) and P19 (end-stage) *Smn^-/-^;SMN2* and wild type (WT) mice. Data are mean ± SEM, n = 3-4 animals per experimental group, two-way ANOVA, Sidak’s multiple comparison test, * *p* < 0.05, ** *p* < 0.01, *** *p* < 0.001, **** *p* < 0.0001. **h-i**. Quantification of NF-κB p50/actin protein levels in the TA of pre-symptomatic (P2) (**h**) and late-symptomatic (P7) (**i**) *Smn^-/-^;SMN2* mice and age-matched WT animals. Images are representative immunoblots. Data are mean ± SEM, n = 3-4 animals per experimental group, unpaired *t* test, ns = not significant (**h**), *p* = 0.0215 (**i**). **j.** qPCR analysis *NF-κB inducing kinase* (*NIK*) in TA muscle of late-symptomatic P7 *Smn^-/-^;SMN2* and age-matched WT animals. Data are mean ± SEM, n = 3-4 animals per experimental group, unpaired *t* test, *p* = 0.0094. **k-q**. qPCR analysis of *parvalbumin* (**k**), *Tweak* (**l**), *Fn14* (**m**), *Pgc-1α* (**n**), *Mef2d* (**o**), *Glut-4* (**p**) and *HKII* (**q**) in TA muscles from P0 (birth), P2 (pre-symptomatic), P4 (pre-symptomatic), P11 (early symptomatic) and P19 (end-stage) *Smn^2B/-^* and WT mice. Data are mean ± SEM, n = 3-4 animals per experimental group, two-way ANOVA, Sidak’s multiple comparison test, * *p* < 0.05, ** *p* < 0.01, *** *p* < 0.001, **** *p* < 0.0001.

Fc-TWEAK was administered by subcutaneous injections using a sterile 0.1 cc insulin syringe at various doses (7.9 μg, 15.8 μg or 31.6 μg) and at a volume of 20 μl either daily, every other day or every four days. Mouse Fc-TWEAK, a fusion protein with the murine IgG2a Fc region, and Ig isotope control were kindly provided by Linda C. Burkly (Biogen) [43].

For survival studies, mice were weighed and monitored daily and culled upon reaching their defined humane endpoint.

For all experiments, litters were randomly assigned at birth and whole litters composed of both sexes were used. Sample sizes were determined based on similar studies with SMA mice.

To reduce the total number of mice used, the fast-twitch tibialis anterior (TA) and triceps muscles from the same mice were used interchangeably for respective molecular and histological analyses.

### Sciatic nerve crush

For nerve crush experiments, post-natal day (P) 7 wild-type (WT) FVB/N mice [39] were anesthetized with 2% isoflurane/oxygen before one of their lateral thighs was shaved and a 1 cm incision in the skin was made over the lateral femur. The muscle layers were split with blunt scissors, the sciatic nerve localized and crushed with tweezers for 15 seconds. The skin incision was closed with surgical glue and animals allowed to recover on a warming blanket. Ipsilateral and contralateral TA muscles were harvested at P14 and either fixed in 4% paraformaldehyde (PFA) for 24 hours for histological analyses or snap frozen for molecular analyses.

### Cardiotoxin injections

Cardiotoxin γ (Cytotoxin I, Latoxan, L8102, Portes les Valence) was dissolved in 0.9% saline and injected 4 µl/g per total mouse weight of a 10 µM solution into the left TA muscle of WT FVB/N mice [39] at post-natal day (P) 10. The right TA was injected with equal volumes of 0.9% saline. During the injection, mice were anesthetized with 2% isoflurane/oxygen and all injections were done using a sterile 0.3 cc insulin syringe. TA muscles were harvested 6 days later and either fixed in 4% PFA for 24 hours for histological analyses or snap frozen for molecular analyses.

### Laminin staining of skeletal muscle

TA muscles were fixed in PFA overnight. Tissues were sectioned (13 μm) and incubated in blocking buffer for 2 hours (0.3% Triton-X, 20% fetal bovine serum (FBS) and 20% normal goat serum in PBS). After blocking, tissues were stained overnight at 4°C with rat anti-laminin (1:1000, Sigma L0663) in blocking buffer. The next day, tissues were washed in PBS and probed using a goat-anti-rat IgG 488 secondary antibody (1:500, Invitrogen A-11006) for one hour. PBS-washed tissues were mounted in Fluoromount-G (Southern Biotech). Images were taken with a DM IRB microscope (Leica) with a 20X objective.

Quantitative assays were performed blinded on 3-5 mice for each group and five sections per mouse. The area of muscle fiber within designated regions of the TA muscle sections was measured using Fiji (ImageJ) [45].

### Hematoxylin and eosin staining of skeletal muscle

TA muscles were fixated in 4% PFA and imbedded into paraffin blocks. For staining, muscles were sectioned (13 μm) and deparaffinized in xylene and then fixed in 100% ethanol. Following a rinse in water, samples were stained in hematoxylin (Fisher) for 3 minutes, rinsed in water, dipped 40 times in a solution of 0.02% HCl in 70% ethanol and rinsed in water again. The sections were next stained in a 1% eosin solution (BDH) for 1 minute, dehydrated in ethanol, cleared in xylene, and mounted with Fluoromount-G (Southern Biotech). Images were taken with a DM IRB microscope (Leica) with a 20X objective. Quantitative assays were performed blinded on 3-5 mice for each group and five sections per mouse. The area of muscle fibre within designated regions of the TA muscle sections was measured using Fiji (ImageJ) [45].

### Cell culture

Both C2C12 myoblasts [46] and NSC-34 neuronal-like cells [47] were maintained in growth media consisting of Dulbecco’s Modified Eagle’s Media (DMEM) supplemented with 10% FBS and 1% Penicillin/Streptomycin (all Life Technologies). Cells were cultured at 37°C with 5% CO_2_. C2C12 myoblasts were differentiated in DMEM containing 2% horse serum for 7 days to form multinucleated myotubes.

Cells were regularly tested for mycoplasma and remained mycoplasma-free.

### *In vitro* siRNA knockdown

For small interfering RNA (siRNA) transfections, C2C12 myoblasts were seeded onto 12-well plates at a 50% confluency and cultured overnight in 2 mL of DMEM. Cells were washed with PBS prior to siRNA transfection, whereby 100 pmol of each siRNA (*Tweak*, *Fn14*, *Smn*) (Invitrogen, assay IDs s233937, s203164, s74017, respectively) in a complex with 10 µl of Lipofectamine RNAi/MAX (Invitrogen) dissolved in OptiMEM solution (Gibco) was added to the cells for three hours. The transfection mix was then substituted either for DMEM without the siRNAs for 1 day or with a differentiation medium mix without the siRNAs for 7 days.

### qPCR

RNA was extracted from tissues and cells either by a RNeasy kit from Qiagen or by guanidinium thiocyantate-acid-phenol-chloroform extraction using TRIzol Reagent (Life Technologies) as per manufacturer’s instructions. The same RNA extraction method was employed for similar experiments and equal RNA amounts were used between samples within the same experiments. cDNA was prepared with the High Capacity cDNA Kit (Life Technologies) according to the manufacturer’s instructions. The cDNA template was amplified on a StepOnePlus Real-Time PCR Thermocycler (Life Technologies) with SYBR Green Mastermix from Applied Biosystems. qPCR data was analyzed using the StepOne Software v2.3 (Applied Biosystems). Primers used for qPCR were obtained from IDT and sequences for primers were either self-designed or ready-made (Supplementary Table 1). Relative gene expression was quantified using the Pfaffl method [48] and primer efficiencies were calculated with the LinRegPCR software. We normalized relative expression level of all tested genes in mouse tissue and cells to *RNA polymerase II polypeptide J* (*PolJ*) [49].

### PCR array

RNA was extracted using the RNeasy® Microarray Tissue Kit (Qiagen). cDNA was generated with the RT^2^ First Strand Kit (Qiagen). qPCRs were performed using RT² Profiler™ PCR Array Mouse Skeletal Muscle: Myogenesis & Myopathy Mouse (PAMM-099Z, SABiosciences) and RT² Profiler™ PCR Array Mouse Glucose Metabolism (PAMM-006Z SABiosciences). The data were analyzed with RT Profiler PCR Array Data Analysis (version 3.5) and mRNA expression was normalized to the two most stably expressed genes between all samples. We used the publicly available database STRING (version 10.5) for network and enrichment analysis of differently expressed genes [50]. The minimum required interaction score was set at 0.4, medium confidence.

### Western blot

Freshly prepared radioimmunoprecipitation (RIPA) buffer was used to homogenize tissue and cells, consisting of 50 mM Tris pH 8.8, 150mM NaCl, 1% NP-40, 0.5% sodium deoxycholate, 0.1% SDS and complete mini-proteinase inhibitors (Roche). Equal amounts of total protein were loaded, as measured by Bradford Assay. Protein samples were first diluted 1:1 with Laemmli sample buffer (Bio-Rad, Hemel Hempstead, UK) containing 5% β-mercaptoethanol (Sigma) and heated at 100°C for 10 minutes. Next, samples were loaded on freshly made 1.5 mm 12% polyacrylamide separating and 5% stacking gel and electrophoresis was performed at 120 V for ∼1.5 hours in running buffer. Subsequently, proteins were transferred from the gel onto to a polyvinylidene fluoride membrane (Merck Millipore) via electroblotting at 120 V for 60 minutes in transfer buffer containing 20% methanol. Membranes were then incubated for 2 hours in Odyssey Blocking Buffer (Licor). The membrane was then probed overnight at 4°C with primary antibodies (P105/p50, 1:1000, Abcam ab32360; Actin, 1:1000, Abcam ab3280) in Odyssey Blocking Buffer and 0.1% Tween-20. The next day, after three 10-minute washing steps with PBS, the membrane was incubated for 1 hour at room temperature with secondary antibodies (goat anti-rabbit IgG 680RD, 1:1000, LI-COR 926-68071; goat anti-mouse IgG 800CW, 1:1000 LI-COR, 926-32210). Lastly, the membrane was washed three times for 10 minutes in PBS and visualized by scanning 700 nm and 800 nm channels on the LI-COR Odyssey CLx infrared imaging system (LI-COR) for 2.5 minutes per channel. The background was subtracted and signal of protein of interest was divided by signal of the housekeeping protein.

### Statistical Analysis

All statistical analyses were done with the most up to date GraphPad Prism software. When appropriate, a Student’s unpaired two-tail *t*-test, a one-way ANOVA or a two-way ANOVA was used. *Post-hoc* analyses used are specified in Figure Legends. Outliers were identified via the Grubbs’ test. For the Kaplan-Meier survival analysis, the log-rank test was used and survival curves were considered significantly different at *p*<0.05.

## RESULTS

### The TWEAK/Fn14 pathway is dysregulated in two SMA mouse models

We firstly investigated the expression of the TWEAK/Fn14 pathway in skeletal muscle of the severe Taiwanese *Smn^-/-^*;*SMN2* mouse model [37], using muscles with reported differential vulnerability to neuromuscular junction (NMJ) denervation (vulnerability: triceps brachii > gastrocnemius > TA > quadriceps femoris) [51]. Muscles were harvested from *Smn^-/-^*;*SMN2* and WT mice at several time points during disease progression: birth (post-natal day (P) 0, pre-symptomatic (P2), early symptomatic (P5), late-symptomatic (P7) and end stage (P10)).

We assessed the expression of *parvalbumin*, a high affinity Ca^2+^-binding protein, which is downregulated in denervated muscle [52, 53] and a marker of muscle atrophy in skeletal muscle of SMA patients and *Smn^−/−^*;*SMN2* mice [54]. We observed a significant decreased expression of *parvalbumin* mRNA during disease progression (Fig. 1a) in SMA mice compared to WT animals, further confirming parvalbumin as a *bona fide* marker of muscle atrophy in SMA [54]. Furthermore, we noted that parvalbumin expression was downregulated at earlier time points in the two most vulnerable muscles (triceps and gastrocnemius) [51] of SMA mice compared to WT animals (Fig. 1a).

We next evaluated the expression of *Tweak* and *Fn14* and observed significant decreased levels of *Tweak* mRNA in muscles of *Smn^-/-^;SMN2* mice during disease progression, except in the quadriceps (Fig. 1b), in accordance with it being a relatively invulnerable SMA muscle [51]. Similarly, we found significantly lower levels of *Fn14* mRNA in all muscles of *Smn^-/-^;SMN2* mice during disease progression (Fig. 1c) compared to WT animals. Interestingly, the decreased expression of *Fn14* in denervated and atrophied muscles of neonatal animals is different to previous reports in adults where denervation-induced atrophy stimulates its expression [26, 27].

As mentioned above, the TWEAK/Fn14 pathway has been reported to negatively regulate the expression of metabolic effectors Klf15, Pgc-1α, Mef2d, Glut-4 and HKII [29]. Given that we have previously published a concordant increased expression of *Klf15* in skeletal muscle of SMA mice during disease progression [55], we next evaluated if the additional downstream metabolic targets were similarly dysregulated in the predicted directions. We indeed observed that the mRNA expression of *Pgc-1α*, *Mef2d*, *Glut-4* and *HKII* was significantly upregulated in muscles of *Smn^-/-^;SMN2* mice at symptomatic time-points (P5-P10) compared to WT animals (Fig. 1d-g), showing an expected opposite pattern to both *Tweak* and *Fn14* (Fig. 1b-c) [29]. Notably, we also found that in most muscles, mRNA levels of *Pgc-1α*, *Mef2d*, *Glut4* and *HKII* were significantly decreased in pre-symptomatic *Smn^-/-^;SMN2* mice (P0-P5) compared to WT animals (Fig. 1d-g), independently of *Tweak* and *Fn14* (Fig. 1b-c).

TWEAK/Fn14 pathway also regulates the canonical and non-canonical NF-κB pathways in skeletal muscle [56, 57]. In pre-symptomatic (P2) TA muscle, we observed no significant difference in the expression of NF-κB1 (p50), a component of the canonical NF-κB pathway, between *Smn^-/-^*;*SMN2* mice and WT animals (Fig. 1h), consistent with normal *Tweak* and *Fn14* levels (Fig. 1b-c). Conversely, there was a significant decreased expression of NF-κB1 (p50) in TA muscle of symptomatic *Smn^-/-^*;*SMN2* mice compared to WT animals at P7 (Fig. 1i), in line with reduced levels of *Tweak* and *Fn14* (Fig. 1b). We also investigated the expression of NF-κB-inducing kinase (NIK), involved in the non-canonical NF-κB activation pathway [58]. We observed that mRNA levels of NIK were significantly increased in TA muscle of P7 *Smn^-/-^*;*SMN2* mice compared to WT animals (Fig. 1j), suggesting that dysregulated activity of the Tweak/Fn14 in skeletal muscle of SMA mice impacts both the canonical and non-canonical NF-κB pathways, which play key regulatory roles in muscle health and metabolism [20, 21].

Finally, we evaluated the expression of the TWEAK/Fn14 signaling cascade in skeletal muscle of the less severe *Smn^2B/-^* mouse model of SMA [38]. TA muscles were harvested from *Smn^2B/-^* mice and age-matched WT animals at P0 (birth), P2 (early pre-symptomatic), P4 (late pre-symptomatic), P11 (early symptomatic) and P19 (end stage). We found a significant decreased expression of *parvalbumin* (Fig. 1k), *Tweak* (Fig. 1l) and *Fn14* (Fig. 1m) in muscle from *Smn^2B/-^* mice during disease progression compared to WT animals, similar to that observed in the more severe *Smn^-/-^;SMN2* SMA mouse model (Fig. 1a-c). We have previously reported the aberrant increased expression of *Klf15* in the TA muscle of *Smn^2B/-^* mice during disease progression [55]. However, we did not observe an increase in expression of *Pgc-1α* (Fig. 1n), *Mef2d* (Fig 1o), *Glut-4* (Fig 1p) and *HKII* (Fig. 1q), suggesting that the negative regulation of these downstream metabolic effectors may be dependent on disease severity, age and/or genetic strain.

We have thus demonstrated that the TWEAK/Fn14 pathway is dysregulated during progressive muscle atrophy in two SMA mouse models.

### Denervation does not affect the Tweak/Fn14 pathway during the early stages of muscle development

As SMA muscle pathology is defined by both intrinsic defects and denervation-induced events, we set out to determine which of these may influence the dysregulation of the Tweak/Fn14 pathway in SMA muscle. We firstly addressed the denervation component by performing nerve crush experiments in which the sciatic nerves of P7 WT mice were crushed and the muscle harvested at P14 [59]. Of note, the sciatic nerve was crushed in only one hindlimb, leaving the other control hindlimb intact. Quantification of myofiber area in TA muscles showed a significant decrease in myofiber size in the nerve crush muscle compared to the control hindlimb (Fig. 2a-c).

**Figure 2.**
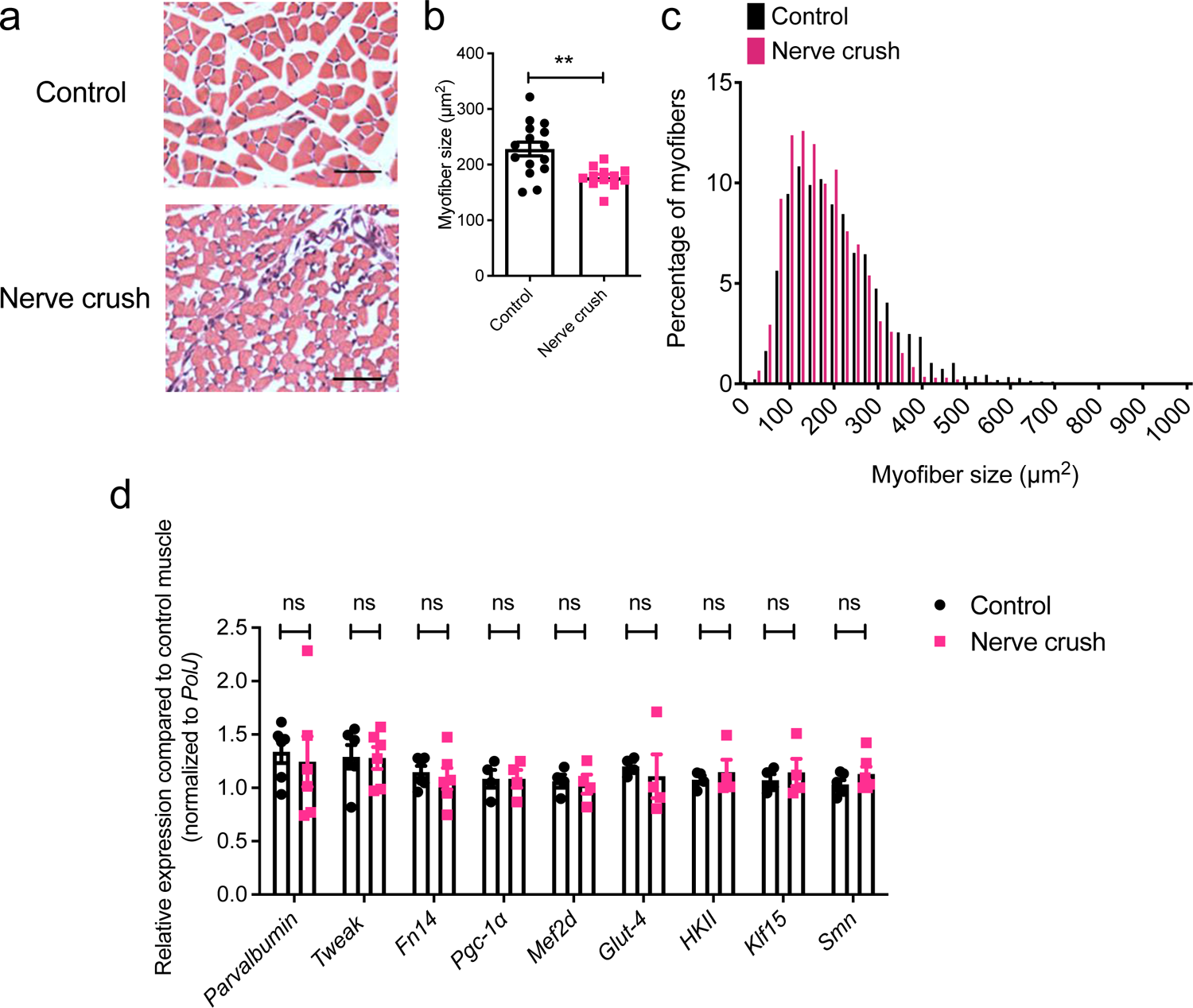
The TWEAK/Fn14 signaling pathway is not dysregulated in denervated muscles of pre-weaned mice. A sciatic nerve crush was performed on post-natal day (P) 7 WT FVB/N mice and both ipsilateral (nerve crush) and contralateral (control) TA muscles were harvested at P14. **a.** Representative images of hematoxylin and eosin-stained cross-sections of control and nerve crush TA muscles. Scale bars = 100 µm. **b.** Myofiber area in control and nerve crush TA muscles. Data are mean ± SEM, n = 3-6 animals per experimental group, unpaired *t* test, *p* = 0.0020. **c.** Myofiber size distribution in control and nerve crush TA muscles. **d.** qPCR analysis of *parvalbumin*, *Tweak*, *Fn14*, *Pgc-1α*, *Mef2d*, *Glut-4*, *HKII*, *Klf15* and *Smn* in control and nerve crush TA muscles. Data are mean ± SEM, n = 4-6 animals per experimental group, two-way ANOVA, uncorrected Fisher’s LSD, ns = not significant.

Expression analyses further revealed that there were no significant changes in mRNA levels of *parvalbumin*, *Tweak*, *Fn14*, *PGC-1α*, *Mefd2*, *Glut-4* and *HKII* in the denervated muscle compared to the control TA muscle (Fig. 2d). Interestingly, while denervation in adult muscle induces a dramatic surge in Fn14 expression [26, 27], this did not occur in the denervated muscles of our pre-weaned mice, suggesting an age and/or development regulatory element to this response. We also investigated the expression of *Klf15* and *Smn* and similarly observed no significant differences between the nerve crush and control muscles (Fig. 2d).

Overall, these results suggest that the dysregulation of parvalbumin and the Tweak/Fn14 pathway in SMA muscle during disease progression is most likely not denervation-dependent.

### Intrinsic muscle injury affects the Tweak/Fn14 pathway during the early stages of muscle development

We next investigated what impact impairing intrinsic muscle integrity would have on the Tweak/Fn14 pathway. To do so, we used cardiotoxin to induce myofiber necrosis. Cardiotoxin was injected in P10 WT mice into the left TA while the right TA was injected with equal volumes of 0.9% saline and used as a control [60]. TAs were harvested after 6 days, a time-point where muscles are still in an immature and regenerating mode [61]. Indeed, analysis of centrally located nuclei showed a significantly increased percentage of regenerating myofibers in cardiotoxin-treated muscles compared to saline-treated TAs (Fig. 3a-b).

**Figure 3.**
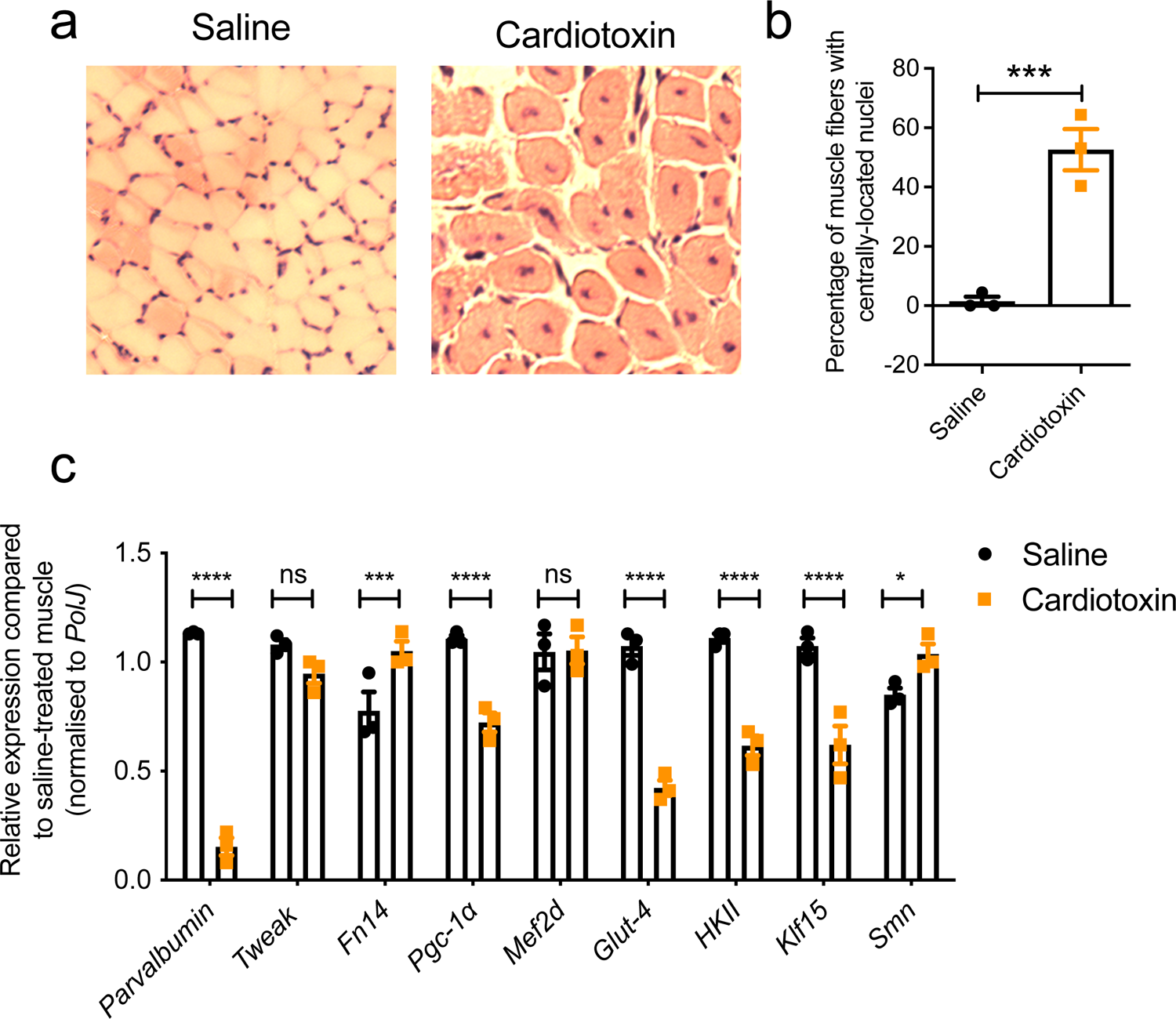
The TWEAK/Fn14 signaling pathway is dysregulated in cardiotoxin-induced muscle necrosis in pre-weaned mice. Cardiotoxin was injected in the left TA muscle of post-natal day (P) 10. The right TA muscle was injected with equal volumes of 0.9% saline. TA muscles were harvested 6 days later. **a.** Representative images of hematoxylin and eosin-stained cross-sections of saline- and cardiotoxin-injected TA muscles. Scale bars = 100 µm. **b.** Percentage of muscle fibers with centrally-located nuclei in saline- and cardiotoxin-injected TA muscles. Data are mean ± SEM, n = 3 animals per experimental group, unpaired *t* test, *p* = 0.0020. **c.** qPCR analysis of *parvalbumin*, *Tweak*, *Fn14*, *Pgc-1α*, *Mef2d*, *Glut-4*, *HKII*, *Klf15* and *Smn* in saline- and cardiotoxin-injected TA muscles. Data are mean ± SEM, n = 3 animals per experimental group, two-way ANOVA, uncorrected Fisher’s LSD, ns = not significant, * *p* < 0.05, *** *p* < 0.001, **** *p* < 0.0001.

We then proceeded with molecular analyses and observed that the atrophy marker *parvalbumin* was significantly downregulated in cardiotoxin-treated TA muscles compared to saline-treated TA muscles (Fig. 3c). *Fn14* mRNA expression was significantly increased after cardiotoxin injury, in accordance with previous research showing that muscle damage conditions activate Fn14 [26]. Conversely, *Pgc-1α*, *Glut-4*, *HKII* and *Klf15* mRNA levels were significantly downregulated (Fig. 3c), supporting their reported negative regulation by the Tweak/Fn14 pathway [29]. Interestingly, *Tweak* mRNA expression remained unchanged, contrary to reports of upregulation following cardiotoxin injury in adult muscle [62], suggesting a differential response in early developmental stages of skeletal muscle. Notably, *Smn* expression was significantly increased in the regenerating muscles compared to saline-treated TA muscles (Fig. 3c), perhaps due to SMN’s role during muscle fiber regeneration [63].

Together, these results demonstrate that intrinsic muscle injury in pre-weaned mice induces a dysregulation of the Tweak/Fn14 signaling cascade. However, the changes were in the opposite direction than that observed in SMA muscles (Fig. 1b), perhaps due to the necrosis and regeneration events that occur following cardiotoxin injury [64], which are not typically found in muscles of SMA mice.

### Genetic interactions between Smn, Tweak and Fn14 in muscle

We next wanted to further understand the potential relationship between dysregulated expression of *Tweak*, *Fn14* and *Smn* in skeletal muscle of SMA mice. To do so, we evaluated the impact of Tweak and Fn14 depletion in the early stages of muscle development by performing molecular analyses on P7 triceps from *Fn14^-/-^* [44], *Tweak^-/-^* [43] and WT mice. In *Tweak ^-/-^* mice, we observed a significant increased expression of *Fn14* with a concomitant significantly decreased expression of *Klf15* compared to WT animals (Fig. 4a). Notably, we found a significant decreased expression of *Smn* in *Tweak^-/-^* triceps compared to WT mice (Fig. 4a), suggesting a direct or indirect positive interaction between Tweak and Smn levels. For their part, *Fn14^-/-^*mice displayed a significant downregulation of *parvalbumin* and a significant upregulation of *Pgc-1α* (Fig. 4b). These analyses further validate the reported negative regulation of Pgc-1α and Klf15 by Fn14 and support the absence of overt pathological muscle phenotypes in young *Tweak^-/-^* and *Fn14^-/-^* mice [26, 65].

**Figure 4.**
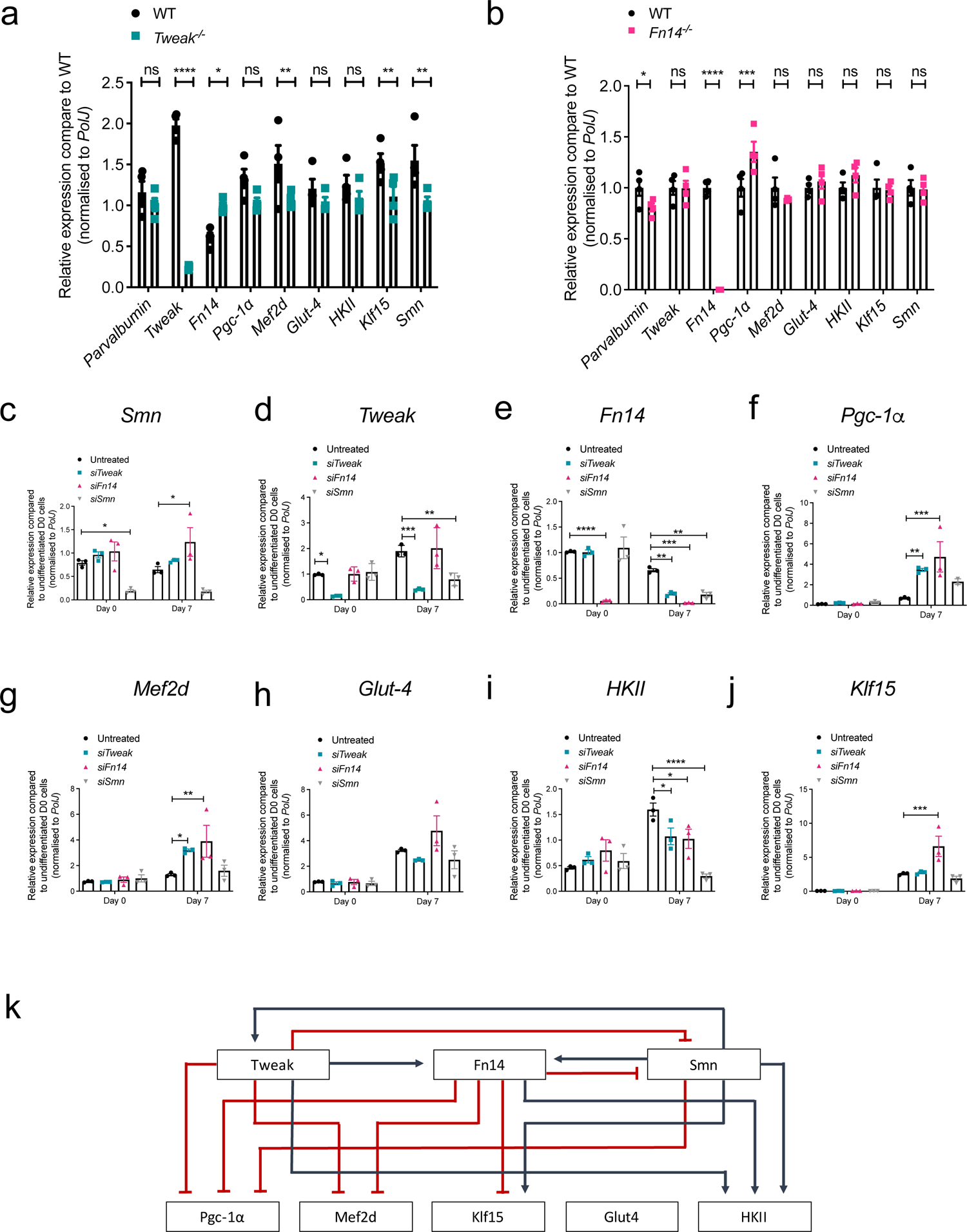
*Smn*, *Tweak* and *Fn14* depletion impact each other’s expression and that of the Tweak/Fn14 signaling pathway. **a-b.** qPCR analysis of *parvalbumin*, *Tweak*, *Fn14*, *Pgc-1α*, *Mef2d*, *Glut-4*, *HKII*, *Klf15* and *Smn* in triceps muscle from post-natal day (P) 7 *Tweak^-/-^* (**a**) and *Fn14^-/-^* (**b**) mice. Data are mean ± SEM, n = 4 animals per experimental group, two-way ANOVA, uncorrected Fisher’s LSD, ns = not significant, * *p* < 0.05, *** *p* < 0.001, **** *p* < 0.0001. **c-j**. qPCR analysis of *Smn* (**c**), *Tweak* (**d**), *Fn14* (**e**), *Pgc-1α* (**f**), *Mef2d* (**g**), *Glut-4* (**h**), *HKII* (**i**) and *Klf15* (**j**) in siRNA-mediated *Tweak*-, *Fn14*- and *Smn*-depleted and control proliferating (Day 0) and differentiated (Day 7) C2C12 cells. Data are mean ± SEM, *n* = 3 per experimental group, two-way ANOVA, Dunnett’s multiple comparisons test, * *p* < 0.05, ** *p* < 0.01, *** *p* < 0.001, **** *p* < 0.0001. **k.** Proposed model of the relationship between Smn and the Tweak/Fn14 signaling pathway. Red lines represent inhibition and blue lines represent activation.

To further dissect the relationship between Smn and the Tweak/Fn14 pathway during myogenic differentiation, we performed siRNA-mediated knockdown of *Smn*, *Tweak* and *Fn14* in C2C12 myoblasts and evaluated the effect on the Tweak/Fn14 signaling in undifferentiated (Day 0) and differentiated (Day 7) cells. Reduced levels of *Smn*, *Tweak* and *Fn14* were significantly maintained in both proliferating and differentiated cells following transfection with *siSmn*, *siTweak* and *siFn14*, respectively (Fig. 4c-e). We observed an interaction between *Smn*, *Tweak* and *Fn14* specifically in differentiated C2C12s, whereby *Smn* expression was significantly upregulated in *Fn14*-depleted D7 cells (Fig. 4c), *Tweak* expression was significantly reduced in *Smn*-depleted D7 cells (Fig. 4d), and *Fn14* levels were significantly decreased in *Tweak*- and *Smn*-depleted D7 cells (Fig. 4e). Similarly, the effects of siRNA-mediated knockdown of *Smn*, *Tweak* and *Fn14* on downstream metabolic effectors were only apparent in differentiated C2C12s (Fig. 4f-j). Indeed, both knockdown of *Tweak* and *Fn14* resulted in a significant upregulation of *Pgc-1α* (Fig. 4f) and *Mef2d* (Fig. 4g). While *Glut-4* expression was neither affected by depletion of *Smn*, *Tweak* or *Fn14* (Fig. 4h), *HKII* mRNA levels were significantly decreased following knockdown of all three (Fig. 4i). Finally, *Klf15* expression was significantly increased in siRNA-mediated knockdown of *Fn14* only (Fig. 4j). The upregulation of *Pgc-1α*, *Mef2d*, and *Klf15* in *Tweak*- and/or *Fn14*-depleted differentiated C2C12 cells is in accordance with the previously reported negative regulation of these genes by the Tweak/Fn14 pathway while the unchanged *Glut-4* and downregulated *HKII* levels were not [18].

Thus, using both *in vivo* and *in vitro* models, we have thus provided evidence for a potential interaction between *Smn*, *Tweak* and *Fn14* and subsequent impact on the Tweak/Fn14 signaling cascade (Fig. 4k). Our results suggest that the aberrant expression of the Tweak/Fn14 pathway in SMA muscle during disease progression may be due to a dynamic interplay between atrophic conditions and the molecular impact, individual and combined, of reduced expression of Smn, Tweak and Fn14 in the early developmental stages of skeletal muscle.

### Overlap of dysregulated myopathy and myogenesis genes and glucose metabolism genes in *SMA*, *Fn14^-/-^* and *Tweak^-/-^* mice

To further decipher the potential contribution(s) of Smn, Tweak and Fn14 depletion to SMA muscle pathology, we used commercially available mouse myopathy and myogenesis qPCR arrays (SABiosciences), which measure expression levels of a subset of 84 genes known to display and/or regulate myopathy and myogenesis. We used triceps (vulnerable) and quadriceps (resistant) from P7 *Smn-/-;SMN2*, *Tweak^-/-^*, *Fn14 ^-/-^* mice. WT FVB/N mice were compared to SMA animals and WT C57BL/6 mice were compared to *Tweak^-/-^* and *Fn14^-/-^* mice to account for differences due to genetic strains. Unsurprisingly, we observed a larger number of significantly dysregulated myopathy and myogenesis genes in triceps of *Smn^-/-^*;*SMN2* mice than in the more resistant quadriceps, some of which overlapped with the subset of genes aberrantly expressed in *Fn14^-/-^* mice and *Tweak^-/-^* mice (Fig. 5a, Table 1, Supplementary File 1). We also used the publicly available database STRING [50] to perform network and enrichment analysis of the shared differently expressed genes in both triceps and quadriceps (Table 1), which revealed that there were no known protein-protein interactions between any of the dysregulated genes and Smn, Fn14 or Tweak (Fig. 5b). Interestingly, the central connectors *Myod1* and *Myf6* were upregulated and *Pax7* was downregulated in the triceps of all three experimental groups (Table 1). Myod1 and Myf6 are key myogenic regulatory factors (MRFs) and are normally upregulated after skeletal muscle injury [66]. Pax7 is a canonical marker for satellite cells, the resident skeletal muscle stem cells [66], and reduced activity of Pax7 leads to cell-cycle arrest of satellite cells and dysregulation of MRFs in skeletal muscle [67]. Furthermore, *Titin* (*Ttn*) was downregulated in the quadriceps muscles of all three mouse models and plays major roles in muscle contraction and force production, highlighted by titin mutations leading to a range of skeletal muscle diseases and phenotypes [68].

**Figure 5.**
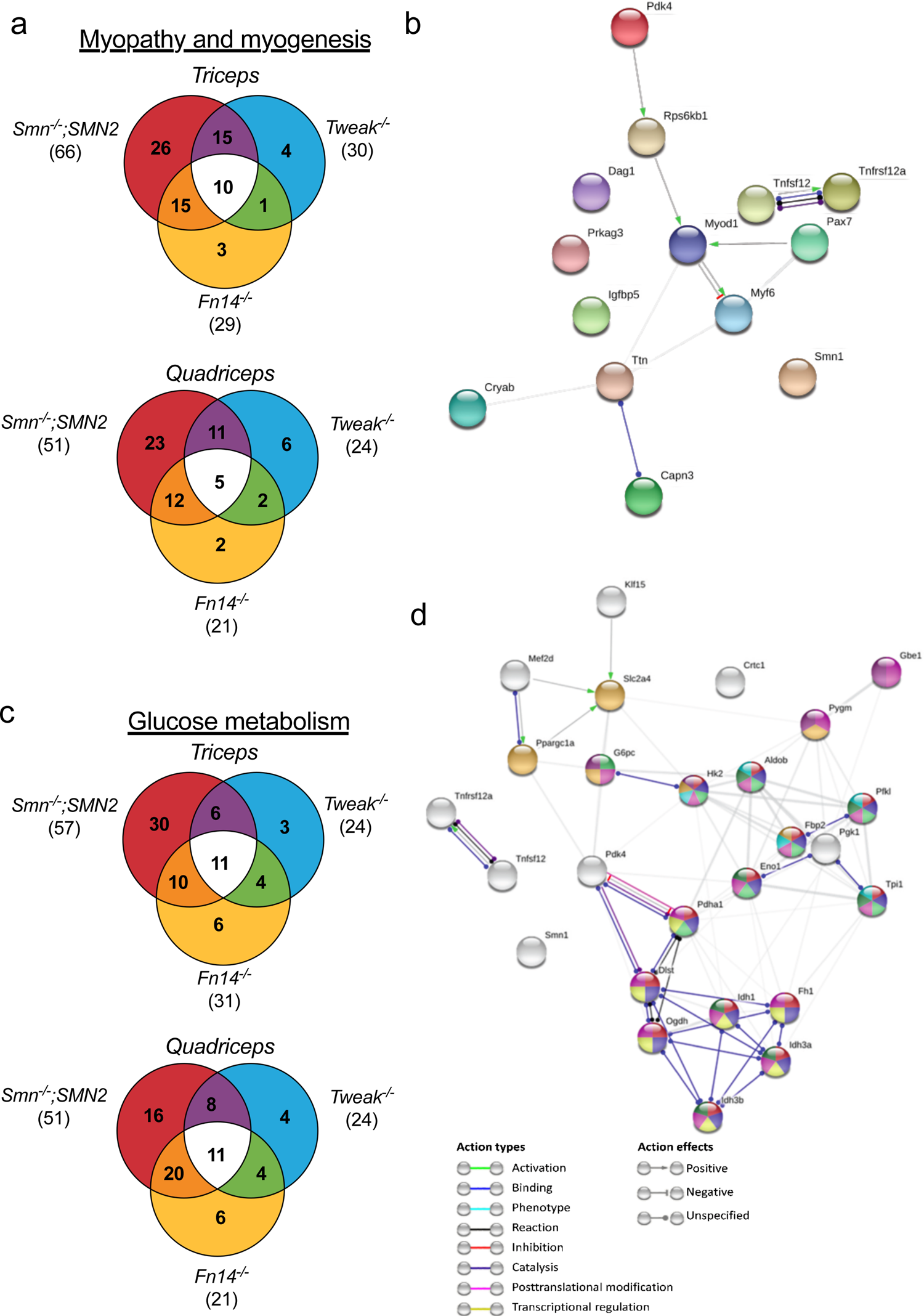
Overlap between dysregulated genes involved in myopathy, myogenesis and glucose metabolism in skeletal muscle of *Smn^-/-^;SMN2*, *Fn14^-/-^* and *Tweak^-/-^* mice. **a.** Venn diagram showing overlap of genes involved in myopathy and myogenesis that are significantly dysregulated in the same direction (either up or downregulated, *p* < 0.05) in triceps and quadriceps muscle from post-natal day (P) 7 compared to *Smn^-/-^;SMN2*, *Fn14^-/-^* and *Tweak^-/-^* mice to age- and genetic strain-matched wild type (WT) mice. **b.** Network and enrichment analysis of the overlap of significantly dysregulated myopathy and myogenesis genes in triceps and/or quadriceps of P7 *Smn^-/-^;SMN2*, *Fn14^-/-^* and *Tweak^-/-^* mice using STRING software. Smn (Smn1), TWEAK (Tnfsf12) and Fn14 (Tnfrsf12a) are included in the analysis. Corresponding protein nodes in the network are highlighted in color. The connection color and shape between proteins represent protein-protein associations (Action types) and if the association is positive, negative or unspecified (Action effects). **c.** Venn diagram showing overlap of genes involved in glucose metabolism that are significantly dysregulated in the same direction (either up or downregulated, *p* < 0.05) in triceps and quadriceps muscle from P7 compared to *Smn^-/-^;SMN2*, *Fn14^-/-^* and *Tweak^-/-^* mice to age- and genetic strain-matched WT mice. **d.** Network and enrichment analysis of the overlap of significantly dysregulated myopathy and myogenesis genes in triceps and/or quadriceps of P7 *Smn^-/-^;SMN2*, *Fn14^-/-^* and *Tweak^-/-^* mice using STRING software. Smn (Smn1), TWEAK (Tnfsf12), Fn14 (Tnfrsf12a), HKII (Hk2), Glut4 (Slc2a4), Pgc-1α (Ppargc1a), Klf15 and Mef2d are included in the analysis. Corresponding protein KEGG pathways with the six lowest FDRs highlighted in color (see Table 3). The connection color and shape between proteins represent protein-protein associations (Action types) and if the association is positive, negative or unspecified (Action effects).

**Table 1.** Myogenesis and myopathy genes significantly dysregulated in the same direction in triceps and quadriceps of P7 *Smn^-/-^;SMN2*, *Fn14^-/-^* and *Tweak^-/-^* mice when compared to P7 WT mice.

| **Gene** | ***Triceps*** | ***Quadriceps*** |
| --- | --- | --- |
| *Calpain3* (Capn3) | Up | Up |
| *Crystallin Alpha B* (Cryab) | Up | — |
| *Dystroglycan 1* (Dag1) | Down | Down |
| *Insulin Like Growth Factor Binding Protein 5* (Igfbp5) | Down | — |
| *Myogenic Factor 6* (Myf6) | Up | — |
| *Myogenic Differentiation 1* (Myod1) | Up | — |
| *Paired Box 7* (Pax7) | Down | — |
| *Protein Kinase AMP-Activated Non-Catalytic Subunit Gamma 3* (Prkag3) | Down | Down |
| *Pyruvate Dehydrogenase Kinase 4* (Pdk4) | Up | — |
| *Ribosomal Protein S6 Kinase B1* (Rps6kb1) | Down | Down |
| *Titin* (Ttn) | — | Down |

Next, as both SMA and the Tweak/Fn14 pathway have both been associated with glucose metabolism abnormalities [29, 69], we performed similar gene expression analyses with commercially available qPCR arrays (SABiosciences) containing a subset of 84 genes known to display and/or regulate glucose metabolism. We found a similar large number of genes that were dysregulated in both triceps and quadriceps muscles of *Smn^-/-^*;*SMN2* mice, some of which overlapped with those differentially expressed in *Fn14^-/-^* and *Tweak ^-/-^* mice (Fig. 5c, Table 2, Supplementary File 2). STRING network and enrichment analysis [50] revealed that there are no known protein-protein interactions between any of the dysregulated genes and Smn, Fn14 or Tweak. Further analysis of Kyoto Encyclopedia of Genes and Genomes (KEGG) pathways composed of the glucose metabolism genes significantly dysregulated in the same direction in triceps and quadriceps muscles of P7 *Smn^-/-^*;*SMN2, Fn14^-/-^* and *Tweak^-/-^* mice as well as the downstream effectors of the TWEAK/Fn14 pathway studied in this project (Pgc-1α, Mef2d, Glut4, Klf15, and HKII) reveals that many aspects of glucose metabolism, such as insulin signaling, glycolysis are dysregulated in Smn-, Tweak- and Fn14-depleted mice (Table 3).

**Table 2.**
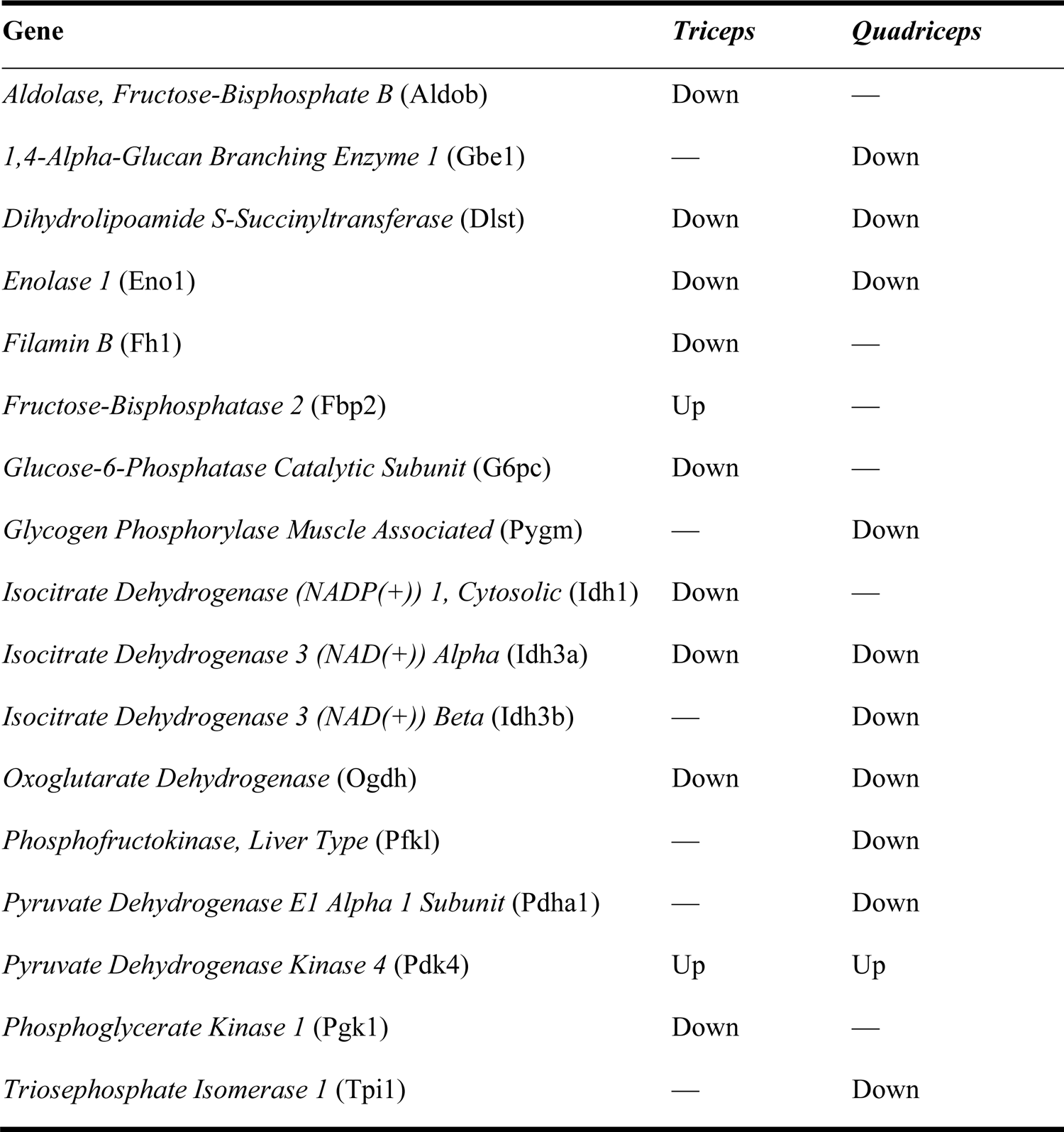
Glucose metabolism genes significantly dysregulated in the same direction in triceps and quadriceps of P7 *Smn^-/-^;SMN2*, *Fn14^-/-^* and *Tweak^-/-^* mice when compared to P7 WT mice.

**Table 3.**
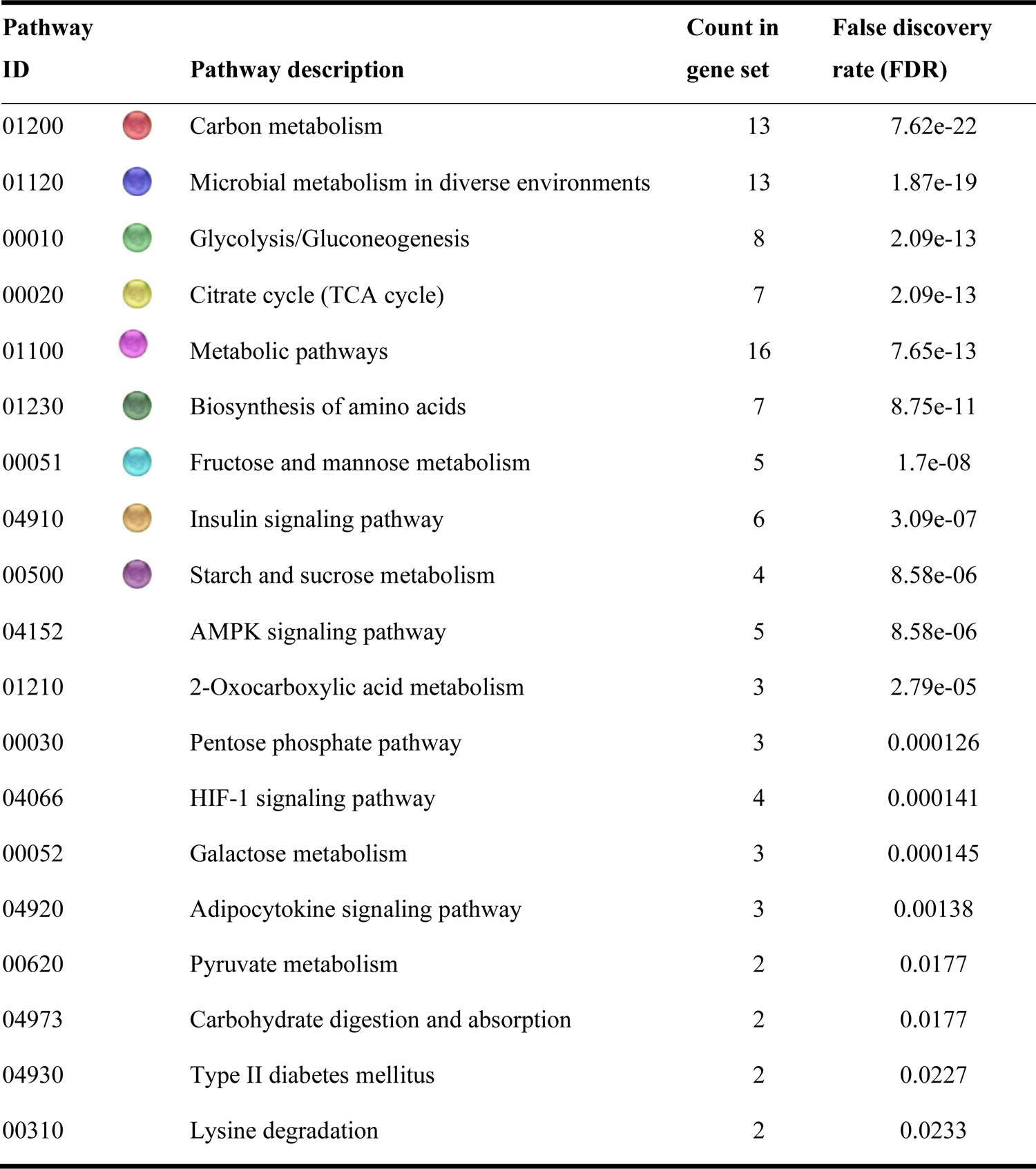
KEGG pathways generated from glucose metabolism genes that were are significantly dysregulated in the same direction in triceps and quadriceps of P7 *Smn^-/-^;SMN2*, *Fn14^-/-^* and *Tweak^-/-^* mice when compared to P7 WT mice.

| Pathway ID |  | Pathway description | Count in gene set | False discovery rate (FDR) |
| --- | --- | --- | --- | --- |
| 01200 |  | Carbon metabolism | 13 | 7.62e-22 |
| 01120 |  | Microbial metabolism in diverse environments | 13 | 1.87e-19 |
| 00010 |  | Glycolysis/Gluconeogenesis | 8 | 2.09e-13 |
| 00020 |  | Citrate cycle (TCA cycle) | 7 | 2.09e-13 |
| 01100 |  | Metabolic pathways | 16 | 7.65e-13 |
| 01230 |  | Biosynthesis of amino acids | 7 | 8.75e-11 |
| 00051 |  | Fructose and mannose metabolism | 5 | 1.7e-08 |
| 04910 |  | Insulin signaling pathway | 6 | 3.09e-07 |
| 00500 |  | Starch and sucrose metabolism | 4 | 8.58e-06 |
| 04152 |  | AMPK signaling pathway | 5 | 8.58e-06 |
| 01210 |  | 2-Oxocarboxylic acid metabolism | 3 | 2.79e-05 |
| 00030 |  | Pentose phosphate pathway | 3 | 0.000126 |
| 04066 |  | HIF-1 signaling pathway | 4 | 0.000141 |
| 00052 |  | Galactose metabolism | 3 | 0.000145 |
| 04920 |  | Adipocytokine signaling pathway | 3 | 0.00138 |
| 00620 |  | Pyruvate metabolism | 2 | 0.0177 |
| 04973 |  | Carbohydrate digestion and absorption | 2 | 0.0177 |
| 04930 |  | Type II diabetes mellitus | 2 | 0.0227 |
| 00310 |  | Lysine degradation | 2 | 0.0233 |

We thus show a shared pattern of aberrantly expressed genes that modulate myogenesis, myopathy and glucose metabolism in SMA, Tweak-depleted and Fn14-depleted skeletal muscle, suggesting that Smn and the Tweak/Fn14 pathway may act synergistically on muscle pathology and metabolism defects in SMA muscle.

### Increasing Tweak activity improves a subset of disease phenotypes in two SMA mouse models

Finally, we evaluated the impact of activating the Tweak/Fn14 pathway on disease progression and muscle pathology in SMA mice. To do so, *Smn^-/-^;SMN2* mice and healthy littermates received a daily subcutaneous injection of Fc-TWEAK (15.8 µg), a fusion protein with the murine IgG2a Fc region [43], starting at birth. We found that Fc-TWEAK did not significantly impact weight or survival of *Smn^-/-^;SMN2* mice compared to untreated and IgG-treated controls (Fig. 6a-b). Additional lower (7.9 µg) and higher doses (23 and 31.6 µg) were also administered but proved to negatively impact weight and survival (Supplementary Fig. 1).

**Figure 6.**
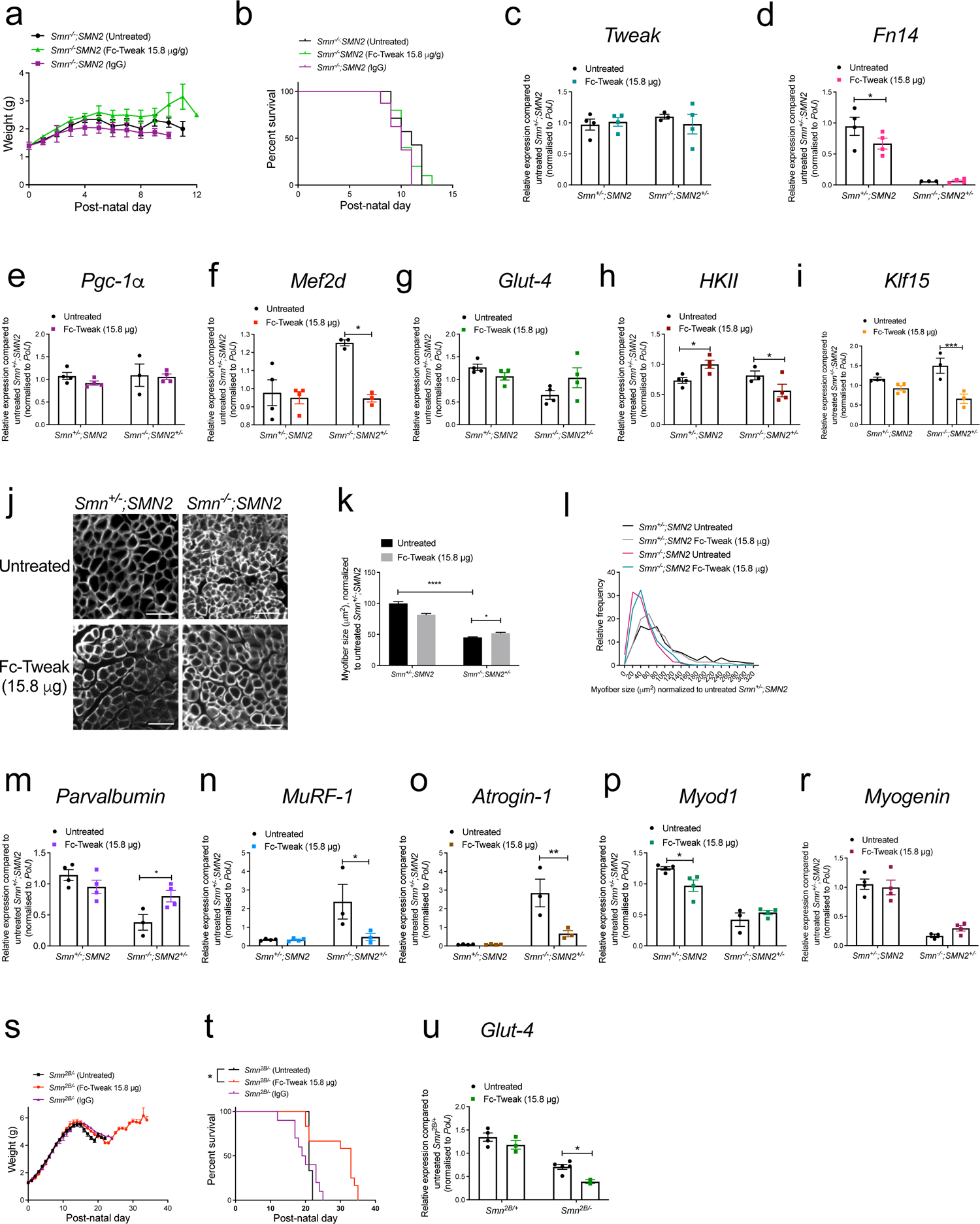
Increasing Tweak activity via Fc-TWEAK improves disease phenotypes in two SMA mouse models. **a.** Daily weights of untreated *Smn^-/-^;SMN2* SMA mice and *Smn^-/-^;SMN2* mice that received daily subcutaneous injections (starting at P0) of Fc-TWEAK or IgG control (15.8 µg). Data are mean ± SEM, n = 7-10 animals per experimental group, two-way ANOVA, Sidak’s multiple comparison test. **b.** Survival curves of untreated *Smn^-/-^;SMN2* SMA mice and *Smn^-/-^;SMN2* that received daily subcutaneous injections of Fc-TWEAK or IgG control (15.8 µg). Data are represented as Kaplan-Meier survival curves, n = 7-10 animals per experimental group, Log-rank (Mantel-Cox). **c-i.** qPCR analysis of *Tweak* (**c**), *Fn14* (**d**), *Pgc-1α* (**e**), *Mef2d* (**f**), *Glut-4* (**g**), *HKII* (**h**) and *Klf15* (**i**) in triceps of post-natal day (P) 7 untreated and Fc-TWEAK-treated (15.8 µg) *Smn^-/-^;SMN2* SMA and *Smn^+/-^;SMN2* health littermates. Data are mean ± SEM, n = 3-4 animals per experimental group, two-way ANOVA, uncorrected Fisher’s LSD, * *p* < 0.05, *** *p* <0.001. **j**. Representative images of laminin-stained cross-sections of TA muscles from P7 untreated and Fc-TWEAK-treated (15.8 µg) *Smn^-/-^;SMN2* SMA and *Smn^+/-^;SMN2* health littermates. Scale bars = 100 µm. **k**. Quantification of myofiber area in the TAs of P7 untreated and Fc-TWEAK-treated (15.8 µg) *Smn^-/-^;SMN2* SMA and *Smn^+/-^;SMN2* health littermates. Data are mean ± SEM, n = 3-4 animals per experimental group (>550 myofibers per experimental group), two-way ANOVA, Tukey’s multiple comparison test, * *p* < 0.05, **** *p* < 0.0001. **l**. Relative frequency distribution of myofiber size in TA muscles of of P7 untreated and Fc-TWEAK-treated (15.8 µg) *Smn^-/-^;SMN2* SMA and *Smn^+/-^;SMN2* health littermates. **m-r**. qPCR analysis of *parvalbumin* (**m**), *MuRF-1* (**n**), *atrogin-1* (**o**), *Myod1* (**p**), and *myogenin* (**r**) in triceps of P7 untreated and Fc-TWEAK-treated (15.8 µg) *Smn^-/-^;SMN2* SMA and *Smn^+/-^;SMN2* health littermates. Data are mean ± SEM, n = 3-4 animals per experimental group, two-way ANOVA, uncorrected Fisher’s LSD, * *p* < 0.05, ** *p* <0.01. **s**. Daily weights of untreated *Smn^2B/-^* SMA mice and *Smn^2B/-^* mice that received subcutaneous injections of Fc-TWEAK or IgG control (15.8 µg) every 4 days (starting at P0). Data are mean ± SEM, n = 9-12 animals per experimental group, two-way ANOVA, Sidak’s multiple comparison test. **t**. Survival curves of untreated *Smn^2B/-^* SMA mice and *Smn^2B/-^* mice that received subcutaneous injections of Fc-TWEAK or IgG control (15.8 µg) every 4 days (starting at P0). Data are Kaplan-Meier survival curves, n = 9-12 animals per experimental group, Log-rank (Mantel-Cox), *p* = 0.0162. u. qPCR analysis of Glut-4 in P15 *Smn^2B/-^* SMA mice and *Smn^2B/-^* mice that received subcutaneous injections of Fc-TWEAK or IgG control (15.8 µg) every 4 days (starting at P0). Data are mean ± SEM, n = 3-4 animals per experimental group, two-way ANOVA, uncorrected Fisher’s LSD, * *p* < 0.05.

Triceps from P7 untreated and Fc-TWEAK-treated (15.8 µg) *Smn^-/-^*;*SMN2* SMA mice and *Smn^+/-^;SMN2* healthy littermates were further processed for molecular analyses of the Tweak/Fn14 pathway. We observed that Fc-TWEAK administration did not influence the expression of *Tweak* (Fig. 6c) or *Fn14* (Fig. 6d) in neither *Smn^+/-^;SMN2* nor *Smn^-/-^*;*SMN2* mice compared to untreated animals. Accordingly, Fc-TWEAK did not induce changes in *Pgc-1α* expression (Fig. 6e). We did observe a significant downregulation of *Mef2d* in Fc-TWEAK-treated muscles of *Smn^-/-^;SMN2* SMA mice compared to untreated animals (Fig. 6f). *Glut-4* mRNA expression remained unchanged in both *Smn^+/-^;SMN2* and *Smn^-/-^*;*SMN2* Fc-TWEAK-treated mice (Fig. 6g). *HKII* was significantly upregulated in muscle of Fc-TWEAK-treated *Smn^+/-^;SMN2* healthy littermates while it was significantly downregulated in Fc-TWEAK-treated *Smn^-/-^;SMN2* SMA mice compared to untreated groups (Fig. 6h). *Klf15* was significantly downregulated in Fc-treated *Smn^-/-^;SMN2* SMA only compared to untreated SMA animals (Fig. 6i). The absence of overt changes in the expression of Tweak, Fn14 and downstream metabolic effectors may be due to the 24 hour time-lapse between the last Fc-Tweak injection and harvest of tissues, which could have led to missing key time-points at which transcriptional profiles were significantly impacted.

Whilst we did not capture the short-term molecular effects of Fc-TWEAK administration, quantification of myofiber area in TA muscles showed that daily Fc-TWEAK treatment significantly increased myofiber area in skeletal muscle of P7 *Smn^-/-^*;*SMN2* mice compared to untreated SMA animals (Fig. 6j-l). Furthermore, the expression of atrophy markers *parvalbumin*, *MuRF-1* and *atrogin-1* [70] was also restored towards normal levels, whereby *parvalbumin* expression was significantly increased (Fig. 6m) whilst *MuRF-1* and *atrogin-1* expression was significantly downregulated (Fig. 6n-o) in triceps of Fc-TWEAK-treated *Smn^-/-^;SMN2* SMA mice compared to untreated SMA animals, further supporting an improvement in muscle health. We did not however detect changes in MRFs *Myod1* and *myogenin* [66] (Fig. 6p-r).

We next assessed the effect of Fc-TWEAK in *Smn^2B/-^* mice, which are typically more responsive to Smn-independent treatment strategies [55,71–73]. Due to the longer treatment period in these mice (20 days) and the observed toxicity in daily injected mice (> 10 days), the *Smn^2B/-^* and *Smn^2B/+^* mice received subcutaneous injections of Fc-TWEAK and IgG control (15.8 µg) every 4 days, starting at birth. Both IgG and Fc-TWEAK did not significantly impact the weight of *Smn^2B/-^* mice compared to untreated SMA animals (Fig. 6s).

However, Fc-TWEAK significantly increased the lifespan of *Smn^2B^*^/−^ mice compared to both IgG-treated and untreated animals (Fig. 6t). Molecular analyses of triceps from P15 animals only showed a significant effect of Fc-TWEAK on the expression of *Glut-4*, whereby it was downregulated in Fc-TWEAK-treated *Smn^2B/-^* mice compared to untreated animals (Fig. 6u). Similarly to above, the limited impact of Fc-TWEAK on the expression of the Tweak/Fn14 signaling cascade is most likely due to the 72-hour time-lapse between the last injection of Fc-Tweak and tissue harvest.

Taken together, our results demonstrate that increasing Tweak activity in SMA mice has the potential to improve weight, survival, and muscle pathology, suggesting that restoring the Tweak/Fn14 pathway in SMA muscle may lead to sustainable therapeutic benefits.

## DISCUSSION

Motor neuron death and muscle pathology bi-directionally impact on each other in SMA. Indeed, while loss of motor neurons significantly contributes to muscle atrophy, there is also evidence for muscle-intrinsic abnormalities in SMA skeletal muscle, which could be directly or indirectly caused by SMN deficiency [6– 8,74,75]. In this study, we addressed the underlying mechanisms of muscle-intrinsic abnormalities leading to muscle pathology in SMA by investigating the role of the TWEAK/Fn14 pathway in muscle atrophy in SMA. To the best of our knowledge, this is the first study to evaluate the TWEAK/Fn14 pathway in SMA and in early stages of muscle development.

Notably, we showed decreased expression of *Tweak* and *Fn14* in skeletal muscle of two distinct SMA mouse models during disease progression, which is contrary to previous reports of increased TWEAK/Fn14 activity in experimental models of atrophy in adult muscle [18,76,77], suggesting that the TWEAK/Fn14 pathway may have distinct roles in skeletal muscle during development and adulthood. Indeed, *Tweak* mRNA expression is significantly lower in skeletal muscle of 30-day-old WT mice compared to 90-day-old animals, suggesting an age-dependent regulation [78]. Moreover, we observed that the dysregulation of the TWEAK/Fn14 pathway in skeletal muscle of pre-weaned mice appears to be influenced by intrinsic myopathy and not denervation, which is in contrast to what has been reported in experimental models of adult muscle denervation [26, 27], further suggesting distinct developmental roles for the Tweak/Fn14 pathway in skeletal muscle. Given that muscles from younger mice are more resistant to surgically-induced denervation than in older mice [79], the TWEAK/Fn14 pathway may contribute to this age-dependent differential vulnerability of muscle to pathological insults. Thus, the role of TWEAK/Fn14 signaling in muscle pathology may be more nuanced and be influenced by a combination of factors such as absolute levels, downstream signaling cascades activated (e.g. canonical vs non-canonical NF-κB signaling pathways), developmental stage of the muscle, state of muscle atrophy (e.g. chronic vs acute) and primary origin of muscle pathology (e.g. denervation vs intrinsic insult) [20, 21].

Another key observation from our study is a potential interaction and/or overlap between Tweak, Fn14 and Smn and their downstream signaling cascades in muscle. It has previously been demonstrated that once Tweak binds to Fn14, the complex will activate several NF-κB molecular effectors, including TRAF6 and IKK [80]. Interestingly, SMN has been reported to prevent the activation of TRAF6 and IKK, thereby negatively regulating the muscle atrophy-inducing canonical NF-κB pathway [81]. These studies thus suggest converging roles for TWEAK, Fn14 and Smn in muscle, which is further supported by our findings. Indeed, we found that independent *Tweak*, *Fn14* and *Smn* depletion had an impact on each other’s expression in differentiated C2C12 cells and murine muscle. Furthermore, there was an overlap of dysregulated myogenesis, myopathy and glucose metabolism genes in SMA, *Fn14^-/-^* and *Tweak^-/-^* mice. Thus, these results suggest that aberrant expression of the TWEAK/Fn14 pathway in SMA muscle may be a consequence of combined events resulting from muscle atrophy events and reduced SMN expression.

In addition, our results in developing mice do support the previously reported negative regulation of the metabolic factors Pgc-1α, Mef2d, Glut4, Klf15, and HKII in adult muscle [29]. Further analyses of a subset of specific glucose metabolism genes showed that about 20% of these genes were dysregulated in the same direction in *Fn14^-/-^*, *TWEAK^-/-^* and SMA mice. Our KEGG analysis of these shared dysregulated metabolic genes further support the potential relationships and roles of TWEAK, Fn14 and SMN involved in the regulation of glucose metabolism. Indeed, the AMPK signaling pathway, found to be aberrantly regulated in *Fn14^-/-^*, *TWEAK^-/-^* and SMA, is as a master regulator of skeletal muscle function and metabolism [82]. Interestingly, a previous study in *SMNΔ7* SMA mice further showed that chronic treatment with the AMPK agonist AICAR prevented skeletal muscle pathology [83]. In addition, AMPK directly phosphorylates PGC-1α [84], which is also dysregulated in *Smn*-, *Tweak*- and *Fn14*-depleted models [85, 86]. We also found that glycolysis and pyruvate metabolic pathways, which culminate in the generation of ATP, are also dysregulated in SMA, *Fn14^-/-^* and *Tweak^-/-^* mice. Interestingly, siRNA-mediated *Smn* knockdown in NSC-34 cells showed a significant decrease in ATP production [87]. ATP was also decreased in *Smn^-/-^*;*SMN2* mice and in Smn morphant zebrafish [88]. These results could explain mitochondrial dysfunction in SMA patients [7]. Thus, our study strengthens the notion of metabolic dysfunctions contributing to SMA muscle pathology and suggests a potential mechanistic link with the TWEAK/Fn14 pathway.

Our findings also confirm that not all skeletal muscles are equally affected in SMA. Indeed, we observed that the SMA skeletal muscle atrophy marker *parvalbumin* was significantly decreased from an earlier timepoint in the vulnerable triceps and gastrocnemius muscles than in the more resistant TA and quadriceps muscles. Notably, we also found that 20% more myogenesis- and myopathy-related genes were dysregulated in the more vulnerable triceps muscles of *Smn^−/−^*;*SMN2* mice compared to the resistant quadriceps muscles. Conversely, the number of glucose metabolism genes dysregulated in SMA triceps and quadriceps muscles was not significantly different. Previous studies have reported that muscle vulnerability is more closely associated with NMJ denervation than with location or fibre type composition [51]. Our results further suggest that denervation events in vulnerable SMA muscles have a more prominent effect on myogenesis and myopathy than on glucose metabolism.

Finally, modulating Tweak activity via Fc-TWEAK in two SMA mouse models led to interesting observations. Firstly, Fc-TWEAK administration specifically increased lifespan in the milder *Smn^2B/-^* mouse model while it did not impact disease progression in the severe *Smn^-/-^*;*SMN2* mice. This is consistent with previous studies, including ours, demonstrating that the *Smn^2B/-^* mice are more responsive to non-SMN treatments, perhaps due to their longer asymptomatic, and therefore adaptable period [55,71–73,89]. At a molecular level, we found that Fc-Tweak differentially impacted the expression of the *Tweak*, *Fn14* and their metabolic effectors in SMA mice and healthy littermates, perhaps reflecting disease-state dependent regulatory mechanisms of the pathway. Importantly, the expression of *Mef2d*, *HKII* and *Klf15* was significantly downregulated in Fc-TWEAK-treated SMA mice, supporting an increased activity of Tweak in the mice and a subsequent restoration towards normal levels of aberrantly regulated Tweak/Fn14 effectors.

As mentioned above, the timing between Fc-Tweak administration and tissue collection may have limited our analysis of the effect of Fc-Tweak on the Tweak/Fn14 signaling cascade. Nevertheless, administration of Fc-Tweak did improve muscle pathology in SMA mice as demonstrated by the partial restoration of molecular markers of muscle health and myofiber size. These results support a role for the TWEAK/Fn14 pathway in maintaining skeletal muscle health and homeostasis [21]. However, it is important to note that the TWEAK/Fn14 pathway is involved in many other tissues and pathologies such as tumor development and metastasis, heart-related diseases [90], kidney injury, cerebral ischemia [91, 92] and autoimmune diseases [93, 94], which could have influenced the overall impact of systemically administered Fc-Tweak on muscle health and disease progression in SMA mice.

## CONCLUSION

In summary, our results demonstrate a potential role and contribution of the TWEAK/Fn14 pathway to myopathy and glucose metabolism perturbations in SMA muscle. Furthermore, our study, combined with previous work in adult models [20, 21], suggests that dysregulation of the TWEAK/Fn14 signaling in muscle appears to be dependent on the origin of the muscle pathology (e.g. denervation vs intrinsic) and developmental stage of skeletal muscle (e.g. newborn, juvenile, adult, aged), further highlighting the differential and conflicting activities of the pathway. Future investigations should be aimed at both furthering our understanding of the relevance of the Tweak/Fn14 pathway in SMA muscle and defining its role in general in maintaining muscle homeostasis throughout the life course.

## LIST OF ABBREVIATIONS

ALS: amyotrophic lateral sclerosis

ANOVA: analysis of variance

cDNA: complementary deoxyribonucleic acid

DEG: differently expressed genes

DMEM: Dulbecco’s Modified Eagle’s Media

FBS: fetal bovine serum

FDR: false discovery rate

GO: gene ontology

H&E: hematoxylin-and-eosin

KEGG: Kyoto Encyclopedia of Genes and Genomes

mRNA: messenger RNA

NF-κB: nuclear factor kappa-light-chain-enhancer of activated B cells

NMJ: neuromuscular junctions

P: postnatal day

*p*: probability value

PBS: phosphate buffered saline

PCR: polymerase chain reaction

PFA: paraformaldehyde

qPCR: quantitative polymerase chain reaction

RIPA: radioimmunoprecipitation

RNA: ribonucleic acid

RNAi: RNA interference

RT-qPCR: reverse transcriptase-quantitative

PCR SEM: standard error of the mean

siRNA: small interfering RNA

SMA: spinal muscular atrophy

STRING: Search Tool for the Retrieval of Interacting Genes/Proteins

TA: tibialis anterior

WT: wild type

## DECLARATIONS

### Consent for publication

Not applicable.

### Availability of data and materials

All data generated or analyzed during this study are included in this published article or in the supplementary information.

## Competing interests

The authors declare they have no competing interests.

## Funding

K.E.M. was funded by the MDUK and SMA Trust (now SMA UK). M.B. was funded by the SMA Trust (now SMA UK) and Muscular Dystrophy Ireland/MRCG-HRB (MRCG-2016-21). S.K. was supported by an ERASMUS grant. P.C. received financial support from the Deutsche Muskelstiftung. R.K. was funded by the Canadian Institutes of Health Research and Muscular Dystrophy Association (USA).

## Authors’ contributions

Conceptualization: M.B.; Methodology: K.E.M, M.B; Validation: K.E.M., M.B.; Formal analysis: K.E.M., E.M., S.K., M.B.; Investigations: K.E.M., E.M., D.A., B.E., S.K., G.H., N.A., M.B.; Writing - original draft preparation: K.E.M, M.B.; Writing – review and editing: K.E.M., E.M., D.A., B.E., S.K., G.H., N.A., P.C., K.E.D., R.K., M.J.A.W., M.B.; Visualization: K.E.M., M.B.; Supervision: P.C., K.E.D., R.K., M.J.A.W., M.B.; Project administration: M.B.; Funding acquisition: R.K., M.J.A.W., M.B.

## Acknowledgements

We would like to thank the staff at the BMS facility at the University of Oxford.

## SUPPLEMENTARY FIGURE LEGENDS

**Supplementary Figure 1.** Effect of varying Fc-TWEAK on disease progression in *Smn^-/-^;SMN2* SMA mice. *Smn^-/-^;SMN2* mice received daily subcutaneous injections of increasing doses of Fc-TWEAK (7.9, 15., 23.7 and 31.6 μg), starting at birth. **a**. Daily weights of untreated *Smn^-/-^;SMN2* SMA mice and *Smn^-/-^;SMN2* mice that received daily subcutaneous injections (starting at P0) of Fc-TWEAK (7.9, 15.8, 23.7 and 31.6 μg). Data are mean ± SEM, n = 5-10 animals per experimental group, two-way ANOVA, Sidak’s multiple comparison test. **b**. Survival curves of untreated *Smn^-/-^;SMN2* SMA mice and *Smn^-/-^;SMN2* mice that received daily subcutaneous injections (starting at P0) of Fc-TWEAK (7.9, 15.8, 23.7 and 31.6 μg). Data are presented as Kaplan-Meier survival curves, n = 5-10 animals per experimental group, Log-rank (Mantel-Cox).

## SUPPLEMENTARY TABLES

**Supplementary Table 1.** Mouse primers used for quantitative real-time PCR.

## SUPPLEMENTARY FILES

**Supplementary File 1.** Myopathy and myogenesis gene expression changes in triceps and quadriceps of post-natal day 7 *Smn^-/-^;SMN2* (SMA), *Tweak^-/-^* (Tweak KO) and *Fn14^-/-^*; (Fn14 KO) compared to age- and genetic strain-matched wild type animals.

**Supplementary File 2.** Glucose metabolism gene expression changes in triceps and quadriceps of post-natal day 7 *Smn^-/-^;SMN2* (SMA), *Tweak^-/-^* (Tweak KO) and *Fn14^-/-^*; (Fn14 KO) compared to age- and genetic strain-matched wild type animals.

## Notes

### Competing Interest Statement

The authors have declared no competing interest.

### Summary of Updates

To add a co-author (RK) who was mistakingly omitted during the submission process but present on the original manuscript document.

