## Supplementary figures and images for "Dysregulation of the Tweak/Fn14 pathway in skeletal muscle of spinal muscular atrophy mice"

### Supplementary Figure 1

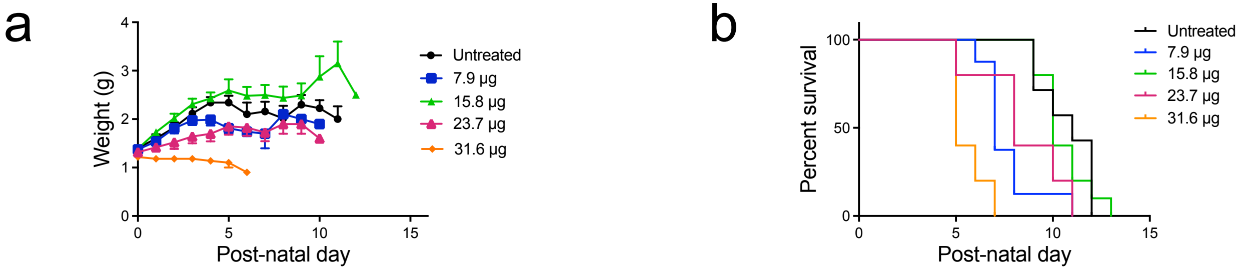
