## Supplementary Table 1 for "Dysregulation of the Tweak/Fn14 pathway in skeletal muscle of spinal muscular atrophy mice"

**Supplementary Table 1. Mouse primers used for quantitative real-time PCR.**

| Mouse | Forward | Reverse |
|---|---|---|
| *Fn14* | 5′-TCGTGTTGGGATTCGGCTTGGT-3' | 5′-ACTTTTCTCTCCGGCGGCATCT-3' |
| *Glut4* | 5′-GACGGACACTCCATCTGTTG-3' | 5′-CATAGCTCATGGCTGGAACC-3' |
| *HkII* | 5′-GAAGGGGCTAGGAGCTACCA-3' | 5′-CTCGGAGCACACGGAAGTT-3' |
| *Klf15* | 5′-TGCGTCGGCACACAGGCGAGAA-3' | 5′-CCGGTGCCTTGACAACTCATCT-3' |
| *Mef2D* | 5′-GCTCCATGCAGTTCAGCAATCCAA-3' | 5′-AGGCTCCATTAGCACTGTTGAGGT-3' |
| *MuRF-1* | 5′-AGGACTCCTGCCGAGTGAC-3' | 5′-TTGTGGCTCAGTTCCTCCTT-3' |
| *MyoD* | 5′-TACAGTGGCGACTCAGATGC-3' | 5′-GAGATGGCGTCCACTATGCT-3' |
| *Myogenin* | 5′-CTACAGGCCTTGCTCAGCTC-3' | 5′-ACGATGGACGTAAGGGAGTG-3' |
| *Parvalbumin* | 5′-GCAAGATTGGGGTTGAAGAA-3' | 5′-GTGTCCGATTGGTACAGCCT-3' |
| *Pgc-1α* | 5′-TGGAGTGACATAGAGTGTGCTGC-3' | 5′-CTCAAATATGTTCGCAGGCTCA-3' |
| *PolJ* | 5′-ACCACACTCTGGGGAACATC-3' | 5′-CTCGCTGATGAGGTCTGTGA-3' |
| *Smn* | 5′-TGCTCCGTGGACCTCATTTCTT-3' | 5′-TGGCTTTCCTGGTCCTAATCCTGA-3' |
| *Tweak* | 5′-AAGTTCACTGAGGGGCCTTGCT-3' | 5′-TGTGAACAAGCTCTGGCTGCCT-3' |
